# Impaired embryo sac cellularization by *PMEI* gene mutation affects gamete specification and twin plants in Arabidopsis

**DOI:** 10.1101/2023.10.17.562779

**Authors:** Isha Sharma, Pinninti Malathi, Ramamurthy Srinivasan, Shripad Ramachandra Bhat, Yelam Sreenivasulu

## Abstract

*Arabidopsis* lines with loss-of-function mutations in the gene encoding **<u>E</u>mbryo sac <u>P</u>ectin <u>M</u>ethyl<u>E</u>sterase <u>I</u>nhibitor** (*Atepmei*) were found to have short silique and high seed sterility. Examination of tissue-cleared mature ovules (FG7-stage) revealed irregularly positioned nuclei within the embryo sacs. Instead of horse-shoe-shaped ovules, defective globular ovules without proper micropylar and chalazal ends were found. Embryo sac cell-type-specific GFP marker expression studies confirmed gamete and accessory cell identity alterations. Egg cell-specific marker (DD45) expression analysis confirmed the presence of multiple egg cells in the mutant embryo sacs, possibly due to defect in embryo sac cellularization. These supernumerary egg cells were functional as evident from the production of twin embryos when supernumerary sperm cells were provided. The results of Ruthenium red and tannic acid-ferric chloride staining of *Atepmei* mutant developing ovules, conferred its interaction with the specific PME in proper cell wall formation and maintenance around embryo sac nuclei which also coincide with its fate as a specific gamete. This is the first report implicating role of cell wall in gamete cell fate determination by altering cell-cell communication. Our analysis of the twin-embryo phenotype of *epmei* mutants also sheds light on the boundary conditions for double fertilization in plant reproduction.

## Introduction

Female gametophyte in angiosperms is a multicellular complex structure deeply embedded into and totally dependent on the mother sporophyte, which considerably limits its investigation. In recent years, significant progress has been made in uncovering details of female gametophyte development through the use of modern microscopy techniques (Susaki et al. 2021) and by devising ingenious molecular and genetic techniques. For instance, a large number of conventional and T-DNA mutants showing defects in female gametophyte development have been generated in *Arabidopsis thaliana*, including those with defects in gamete specification within the embryo sac (Bemer et al., 2008; Portereiko et al., 2006; Steffen et al., 2008). In a typical Polygonum-type embryo sac, eight genetically identical haploid nuclei generated after three rounds of mitosis, migrate to occupy distinct stereotypic positions within a syncytial structure, and cellularize to differentiate into synergids (2), egg cell (1), central cell (1) and antipodal cells (3). How these cell specifications are achieved in a developing female gametophyte is still an important open question in plant reproductive biology. Pagnussat et al. (2009) proposed that auxin gradient within the embryo sac contributes to cell identity specification. However, other studies have failed to find the auxin gradient within the embryo sac (Lituiev et al., 2013). Involvement of *LACHESIS* (*LIS*), *CLOTHO*/*GAMETOPHYTE FACTOR 1* (*CLO*/*GFA1*) and *WYRD* (*WYR*) in the maintenance of gametic versus accessory cell fate has been reported by different researchers Gross-Hardt et al., 2007; Moll et al., 2008). Volz et al. (2012) found *LIS* expression is necessary in the egg cell for the specification and development of synergid cells. Likewise, involvement of type I MADS-domain transcription factors in central cell specification has been reported (Bemer et al., 2008; Portereiko et al., 2006; Steffen et al., 2008). It was demonstrated that diSUMO-like protein (Srilunchang et al., 2010) and a LOB-domain transcription factor (Evans, 2007; Guo et al., 2004) are involved in embryo sac differentiation in maize. Defects in differentiation of gamete cell types both in male and female gametophytes was observed in *RETINOBLASTOMA RELATED* (*RBR*) gene mutant of *Arabidopsis* (Inze and De Veylder, 2006; Johnston et al., 2008; Johnston et al., 2010; Chen et al., 2009). Supernumerary egg cells have been identified as likely because of the trans-differentiation of synergids in *eostre* mutants (Lawit et al., 2013; Pagnussat et al., 2007). In all these mutants including *lis*, *clo*, or *wyr* mutants additional egg cells were identified but their functionality was not demonstrated. Loss of *AMP1* function leads to supernumerary egg cells at the expense of synergids (Kong et al., 2015), enabling the generation of dizygotic twins. Mutant analysis has shown a strict correlation between nuclear position and cell fate (Kong et al., 2015; Grob-hardt et al., 2007; Pagnussat et al, 2007; Moll et al., 2008; Kirioukhova et al., 2011). However, it is still unknown whether nuclear position determines cell fate, as little spatiotemporal information is available on the detailed nuclear dynamics and cell fate specifications. In particular, the roles of embryo sac cellularization and cell wall composition in differentiation of gamete specification have not been investigated so far. Investigation of an *Arabidopsis thaliana* T-DNA promoter trap mutant showing GUS expression in the embryo sac (Sharma et al., 2015) revealed mis-specification of gamete cells leading to high seed sterility. Here we report that the loss-of-function mutation in the *At3g17150* gene annotated to code for a “pectin methylesterase inhibitor” results in this shift in cell fate thereby revealing the role of cell-wall in determining cell fate of female gametophyte and subsequent embryo development.

## Results

### Phenotype of the mutant and transmission of mutant alleles

An *Arabidopsis thaliana* mutant showing high seed sterility was identified in a T-DNA promoter trap mutant population. This line, designated as GUS-650, carries a promoter-less *uidA* reporter gene and showed GUS expression in the ovules (for details see Sharma et al. 2015). Further, the T-DNA was found to be inserted in the 5′ UTR of *At3g17150* gene coding for *<u>P</u>ectin <u>M</u>ethyl<u>E</u>sterase Inhibitor* (*PMEI*) and was specifically expressing in the mature embryo sacs. Therefore, we renamed the mutant as <u>E</u>mbryo-sac-specific *<u>P</u>ectin <u>M</u>ethyl<u>E</u>sterase Inhibitor* (*Atepmei1-1*). To better understand and further confirm the mutant phenotype, another T-DNA insertion line having insertion in the same gene was obtained from ABRC and is named as *Atepmei1-2*. *Atepmei1-2* carries T-DNA in the promoter region of *At3g17150* (Fig. 1). Thus, *Atepmei1-1* is a loss-of-function mutation for *At3g17150* gene. Both the mutants were found to show similar phenotypeand proceeded to study the mutant for *At3g17150*.

**Figure. 1.**
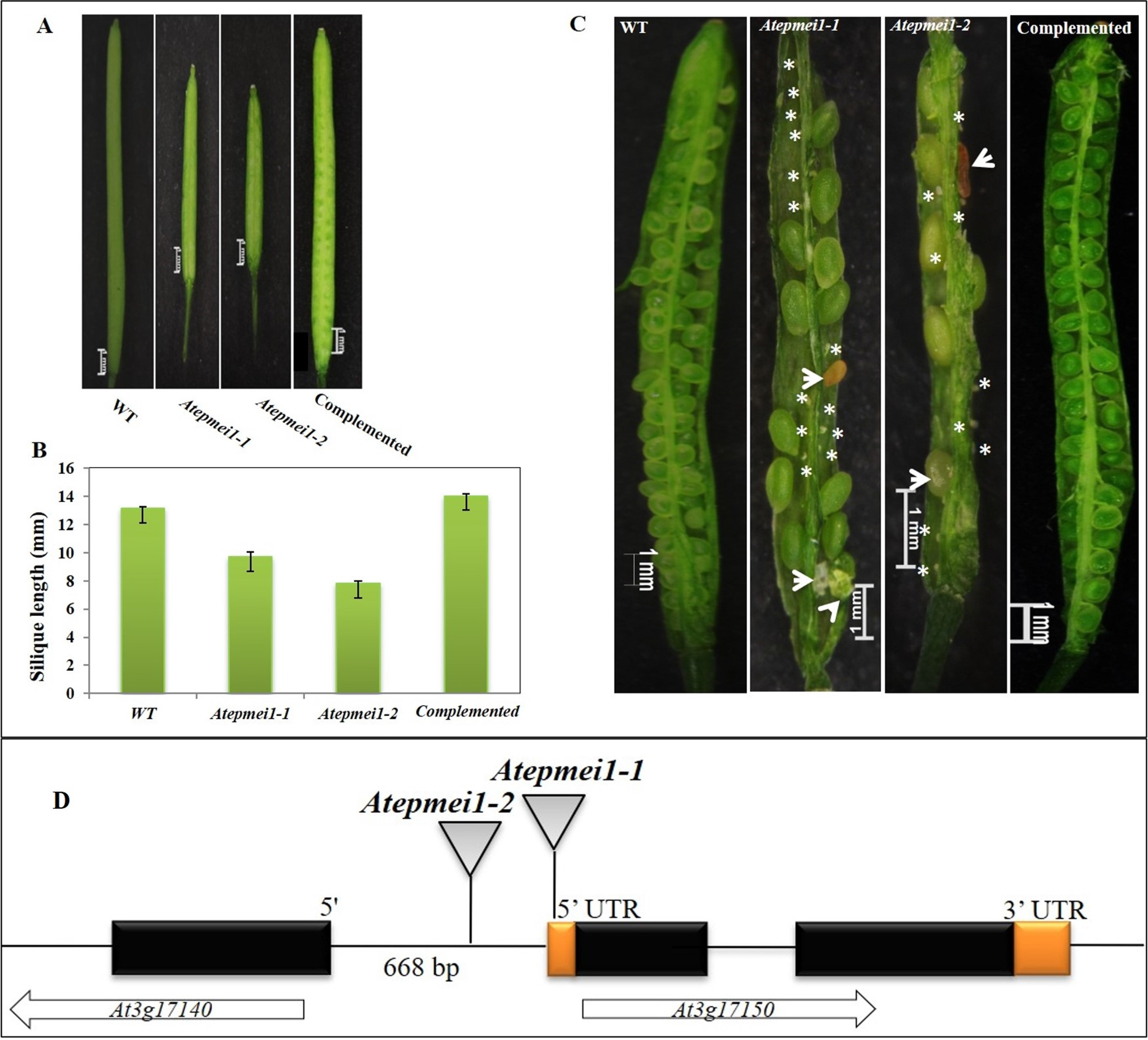
Silique length and seed sterility in *Atepmei* mutant and complemented plants. A) Silique phenotype. B) Bar graph showing silique length (mean ± SE). C) Dissected silique from WT, *Atepmei* mutants, and complemented plants showing seed sterility. Aborted ovules and degenerating seeds in mutant are marked with asterisk and arrowheads, respectively. D) Schematic diagram showing T-DNA insertions in *Atepmei* mutants. T-DNA insertions were in 5’ UTR and in the promoter region in *Atepmeil-1* and *Atepmeil-2,* respectively.

Both the mutants had short siliques as compared with WT (Figure 1) (silique length: WT – 13.11±0.14 mm; *Atepmei1-1 –* 9.68±0.35 mm and *Atepmei1-2 –* 7.79±0.15 mm). Homozygous mutant plants showed 52.82±0.78% and 45.97±1.34 seed sterility in *Atepmei1-1* (N=2444) and *Atepmei1-2* (N=1231), respectively, as compared with WT which showed only 2.78±0.34% seed sterility. Distorted segregation ratio analysis was employed to identify differences in transmission frequency of mutant alleles from the male and the female side as a function of Kanamycin gene transmission present in the T-DNA. For this, reciprocal crosses were performed between the mutant and WT. Analysis of the data showed that the transmission efficiency (TE) through the female gamete is 45.72% and 56.75% in *Atepmei1-1* and *Atepmei1-2*, respectively. Whereas TE through the male gamete was 97.43% and 100% in *Atepmei1-1* and *Atepmei1-2* mutants, respectively (Table S1). In the reciprocal cross, it was observed that the T_1_ seedlings of both mutant lines showed the kanamycin resistant (Kan^r^) to sensitive (Kan^s^) ratio of 0.45:1 and 0.56:1, respectively. On the other hand, in the reciprocal cross, where WT was used as the female and heterozygous mutant plants as the male parent, exhibited Kan^r^:Kan^s^ ratio of ∼1:1 in both the mutant lines.

We examined tissue-cleared anthers under the Differential Interference Contrast (DIC) microscope to see deviations, if any, in male gametophyte development in these mutant lines. As expected, no morphological abnormalities were observed in pollen grains (Upper panel: Supplementary Figure 1). Further, with Alexander’s staining, pollen of *Atepmei* mutants stained pink just like the WT pollen, which indicated their viability (Lower panel: Supplementary Figure 1). These results clearly show that the mutant allele is normally transmitted from the male side but causes defects in the female side. However, the mutation is not completely lethal and yields homozygous mutant plants that survive and perpetuate albeit with high seed sterility.

Homozygous *Atepmei1-1* mutant plants were transformed with p35S::*AtEPMEI* (cDNA) gene cassette for complementation. Both silique length and seed set were restored in the complemented (T_3_ generation) plants. Siliques of the complemented plants showed 98.74% seed set (Figure 1C). All the selected complemented lines showed reverted mutant phenotypes i.e., WT-like silique length, seed set, and normal seed development (Supplementary Figure 2). These results confirmed that *Atepmei1* is a loss-of-function mutation for the *At3g17150* gene and the observed phenotypes are due to loss of this gene function (Figure 1A-C).

### Analysis of female gametophyte development

To identify the nature of seed sterility, tissue-cleared mature ovules from both the mutants were examined at different stages of female gametophyte development using DIC microscope. Ovule development proceeded normally till eight-nucleate embryo sac stage. Mature ovule from WT at the FG7 stage revealed all eight nuclei precisely positioned within the embryo sac i.e., synergids and egg cell at the micropylar end, central cell in the middle and antipodals at the chalazal end of the embryo sac (Fig. 2A). However, mature pre-fertilized ovules from both mutants displayed abnormally positioned nuclei at the micropylar end of the embryo sac (Figure 2B-I). In some of the ovules, the nuclei were arranged erratically (Fig. 2B, C) whereas in others they appeared fused at the micropylar end, and were unable to differentiate into the egg and synergid cells (Figure. 2D, E). Ovules with persistent antipodals at FG7 stage, were also found in some ovule (Figure 2B, D, E). In some others, randomly distributed nuclei with their own cell walls were observed within the mutant embryo sacs (Figure 2F, G, H). Also, ovules with distorted morphology lacking distinction between the micropylar and the chalazal ends were also observed (Figure 2I).

**Figure. 2:**
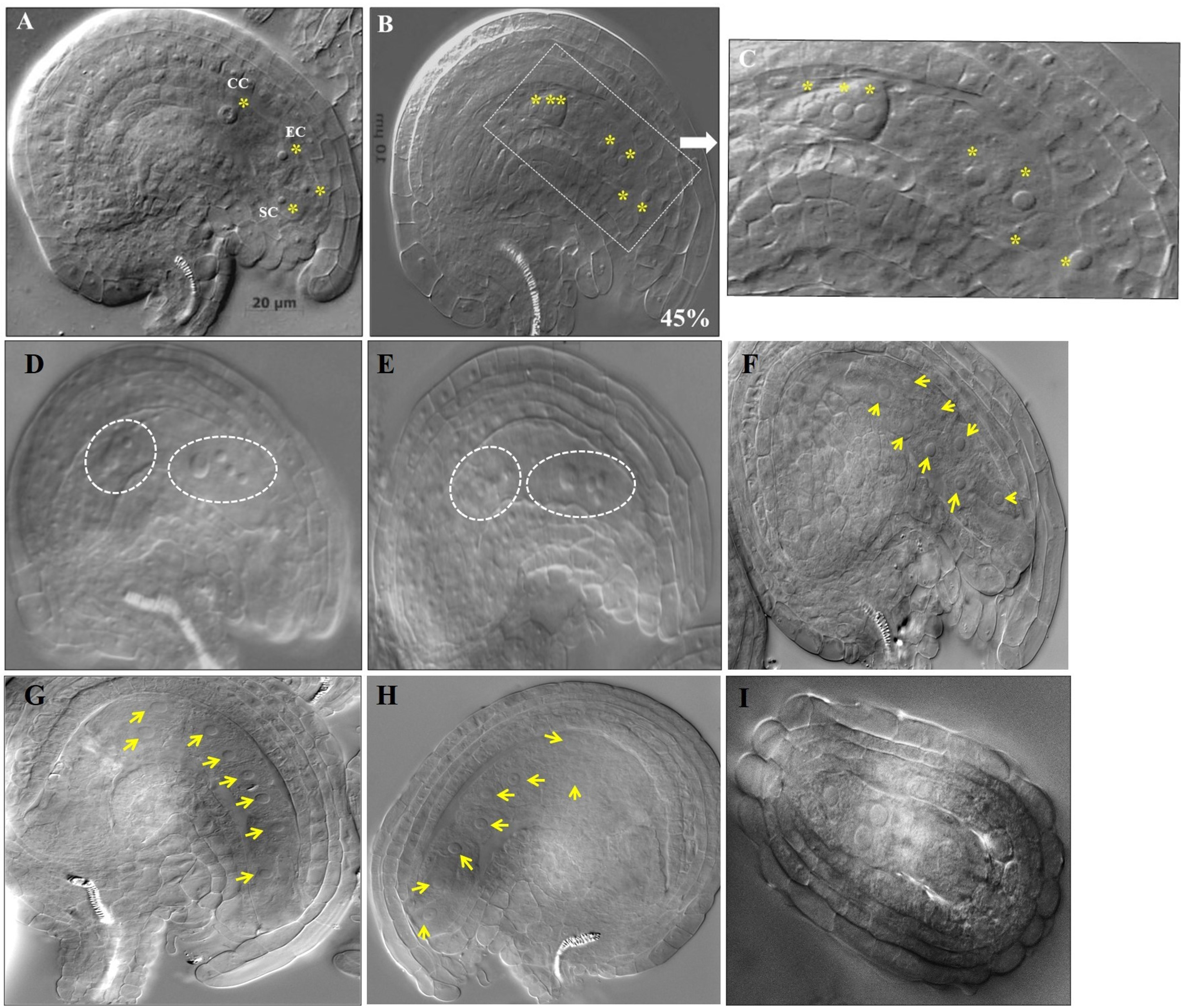
Phenotypic characterization of abnormalities in pre-fertilized female gametophytes of *Atepmeil* homozygous mutant plants. A) Wild-type mature ovule. B-H) *Atepmei* mutant mature ovule. C) Closeup view of B showing linearly arranged nuclei in the mutant embryo sac. D-E) Abnormally positioned nuclei in *Atepmeil-1* and *Atepmeil-2* mutants, respectively. Nuclei are marked with white circles. F-H) Nuclei arranged in a row with wall like structures around them. Nuclei are marked with arrows.

Post-fertilization, mutant ovules displayed zygote-like structures/embryos lying at different locations within the abnormally shaped embryo sacs (Fig. 3). In addition, globular-like embryos were seen in which the suspensor end failed to differentiate properly (Fig. 3B, C, E, F). Further, developing embryos were found mislocated i.e., present elsewhere than at the micropylar end (Supplementary Figure 3). A significant proportion of ovules (19.82% and 17.13% in *Atepmei1-1* and *Atepmei1-2,* respectively) appeared aborted with degenerating embryo sacs and some lacked nuclei. It was also observed that persistent antipodals were prominent in many of the pre-fertilized mutant ovules (Figure 2B-E). Nevertheless, normal WT-like embryo sacs with properly positioned nuclei were also observed (31.19% and 30.27% in *Atepmei1-1* and *Atepmei1-2*, respectively). Interestingly, twin embryos without endosperm were also observed in ∼0.4% of the embryo sacs (Figure 4).

**Figure. 3.**
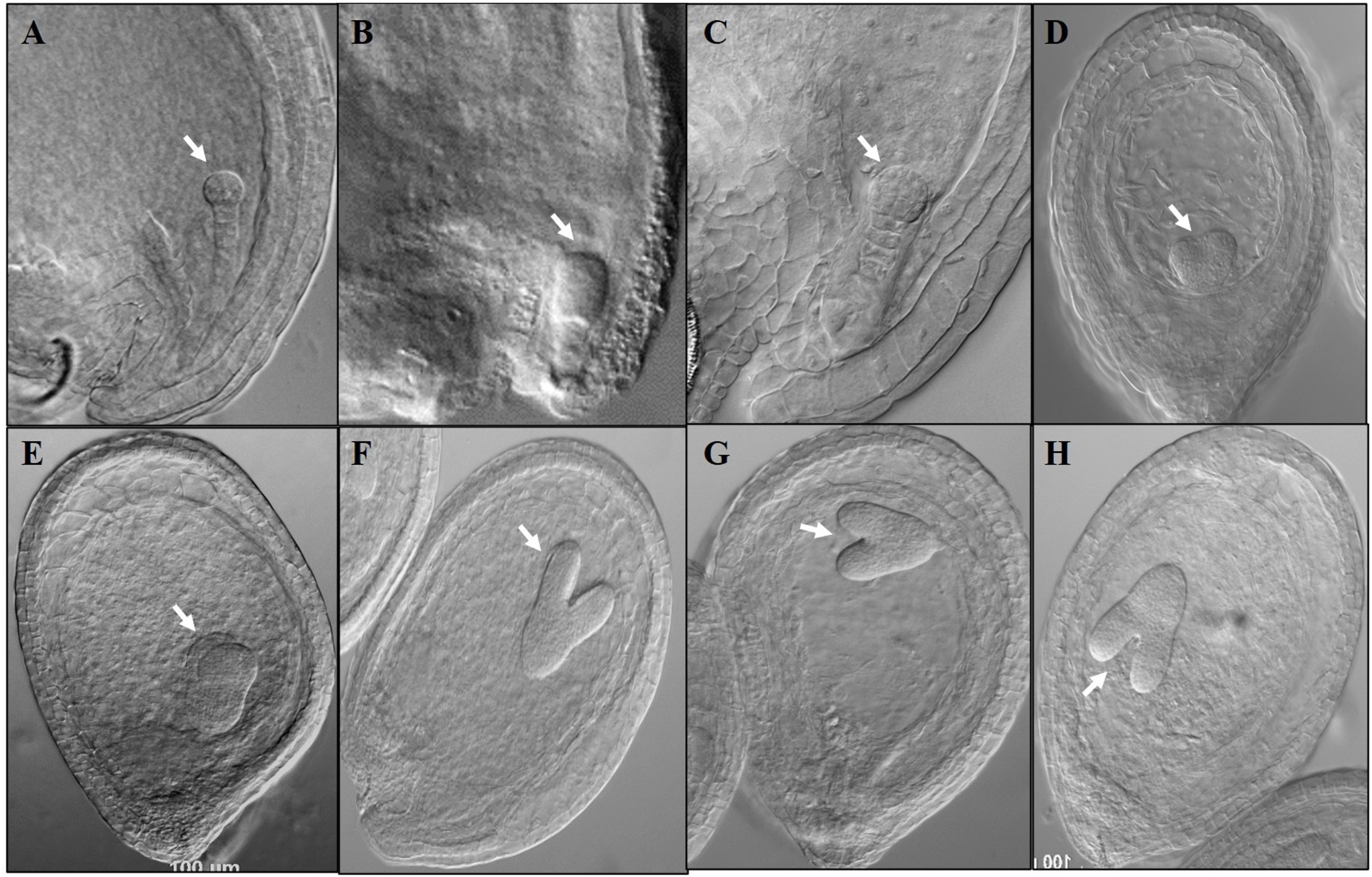
Abnormalities in post-fertilized female gametophytes of Atepmei homozygous mutant plants. A) Wild-type ovule showing globular embryo with suspensor. B) and C) Defective embryos in Atepmeil-1 and Atepmeil-2, respectively. D and E) Abnormal embryo sac and defective embryos in Atepmeil-1 and Atepmeil-2, respectively. F-H) Abnormal embryo sacs with growing embryos at different locations. Arrows indicate the embryos.

**Figure 4.**
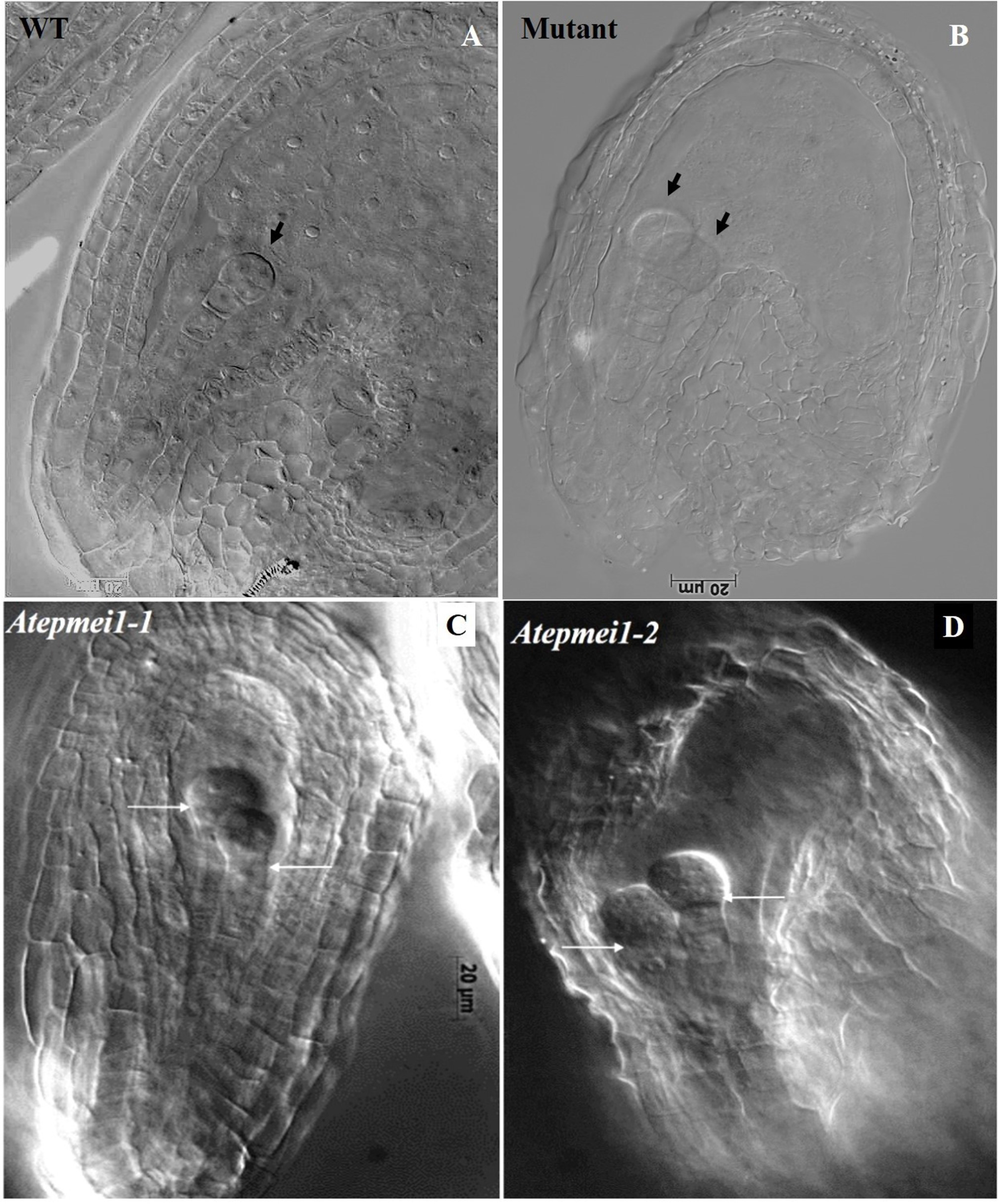
Post-fertilized ovule phenotypes depicting two embryos in *Atepmeil* mutant ovules. A) Single embryo in WT ovule, B-C) Twin embryos in selfed *Atepmeil-1,* and *Atepmeil-2,* respectively, D) *Atepmeil-2* fertilized with WT pollen. Embryos were marked with arrows.

Different categories of embryo sacs were scored and their frequencies are compiled in Table 1. In the majority of ovules (48.97% and 52.58% in *Atepmei1-1* and *Atepmei1-2,* respectively), the nuclei in the embryo sac were arranged in a row or appeared fused at the micropylar end which could not be categorized as either egg cell or synergid cell (Figure 2).

To find whether fertilization with WT pollen would affect embryo development, homozygous mutant plants were crossed with WT pollen and the ovules were examined four days after pollination (4 DAP). In 13.94% crossed seeds (total number of seeds observed = 208) of *Atepmei1-1* and 7.54% crossed seeds (total number of seeds observed =159) of *Atepmei1-2* twin embryos were seen (Figure 4). In the reciprocal cross (WT as female and mutant as male) no twin embryos were observed (total number of seeds observed = 200), which indicated that twin embryos arise due to defect in female gamete development and e*pmei* mutant ovules have more than one egg cell (supernumerary egg cell) which are functional giving rise to twin embryos. In every case, ovules with twin embryos did not progress beyond globular shaped embryo stage. Further, no twin seedlings were observed in the progenies of homozygous mutant plants indicating that the ovules with twin embryos do not progress to maturity. Ovules with twin embryos lacked endosperm and this perhaps led to the arrest of embryos at the globular stage. Since each pollen carries only two sperm nuclei, and if they fertilize the normal and the supernumerary egg cells then the central cell will remain unfertilized. Such unfertilized central cell will fail to differentiate into endosperm. Thus, lack of central cell fertilization appears to be the reason for non-recovery of twin seedlings in the progenies of mutants.

### Fertilization of *pmei1* ovules with *tetraspore* (*tes*) pollen led to development of seeds with twin embryos

To assess whether supplying additional male gametes to fertilize multiple egg cells and also the central cell would lead to development of seeds with twin embryos and endosperm, mutant plants were crossed with pollen from *tetraspore* (*tes*) mutant. Pollen of the *tes* mutant often contain more than two sperm cells. Pollination of *pmei* with *tes* pollen strongly decreased the percentage of abortive seeds from 52.8% (mutant selfed) to 21.5% (mutant crossed with *tes*) and restored endosperm development in ovules containing twin embryos (Figure 5B). Further, twin embryos progressed through torpedo and bent cotyledon stage and developed into mature seeds, which subsequently germinated to give rise to twin seedlings and adult plants (Figure 5D, E, H). These twin plants were further confirmed for the presence of *epmei* and *tes* mutant backgrounds by PCR with insertion-specific forward primer and T-DNA-specific reverse primer for the former gene and gene-specific forward and T-DNA-specific reverse primer for the later gene (Fig. 5G). These primer combinations amplified the expected sizes of the amplicons for both the genes i.e., 600 bp (*epmei*) and 4 kb (*tes*) which endorses the mutant background in these plants. These results also confirmed that *Atepmei* mutants produce multiple egg cells and they are fully functional.

**Figure 5.**
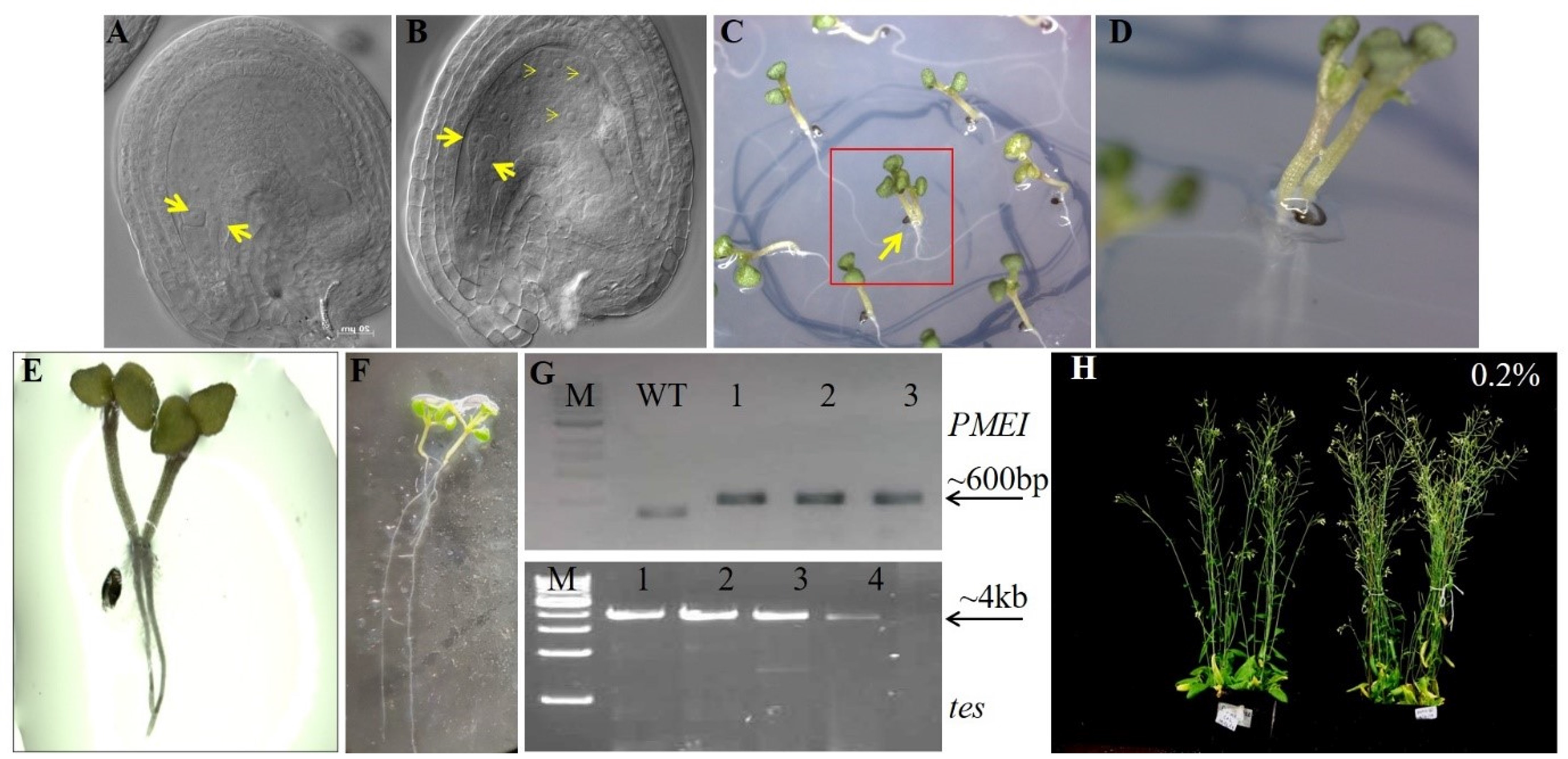
Twin embryos, twin seedlings and twin plants obtained after pollination of *Atepmei* ovules with *tes-4* pollen. **A)** Ovule of self-pollinated *Atepmei* mutant, B) Ovule of *Atepmei* (♀) ♂ *tes-4* (cT) showing proper development of twin embryos and endosperm. C) Germinated F_1_ single and twin seedlings (within rectangle) from *Atepmei* (♀) ♂ *tes-4* (cT). D) Closer view of twin seedlings shown in “C”. E - F) Germinated seedlings showing normal shoot and root development. G) Confirmation of hybridity of twin seedlings through PCR amplification of parental *Atepmei* and *tes-4* alleles, H) Adult twin plants.

The facts that in the mutant, the distribution of the nuclei within the embryo sac was often random (Figure 2) and the developing embryos were found at locations other than the micropylar region (Figure 3) suggested that the egg cell specification was not based on its location within the embryo sac. Therefore, we decided to check the cell specification and their location within the embryo sacs of *Atepmei* mutants. For this, homozygous *Atepmei1-1* and *Atepmei1-2* lines were separately crossed with different cell-specific marker lines i.e., DD31::GFP (synergid cell-specific), DD45::GFP (egg cell-specific), DD65::GFP (central cell-specific) and DD1::GFP (antipodal cell-specific) marker lines. The F_1_ plants were examined for GFP expression in the embryo sacs.

In WT ovules, egg cell-specific marker (DD45::GFP) shows GFP expression at a single position i.e., in-between the central cell and synergid cells (Figure 6A). In *Atepmei1-1* and *Atepmei1-2* mutant ovules, 44-59% embryo sacs showed no GFP expression (Supplementary Table 2) and 18-24% ovules showed normal WT-like GFP fluorescence. Some ovules from the mutant plants showed GFP fluorescence in abnormal positions. In 13.96% of ovules, GFP fluorescence was documented at two positions, one at the normal egg cell position and the other at a synergid position (Figure 6B). In 7.32 -17.33% of ovules diffused GFP fluorescence was documented throughout the embryo sac (Figure 6C). Further, in 13.97% of ovules, three GFP spots were documented, two spots at the micropylar end, and one at the chalazal end (Figure 6D). GFP fluorescence was also observed in the chalazal end at the antipodal position (Figure 6E). GFP fluorescence was also found at the synergids cell position at the micropylar end (Fig 6F).

**Figure 6.**
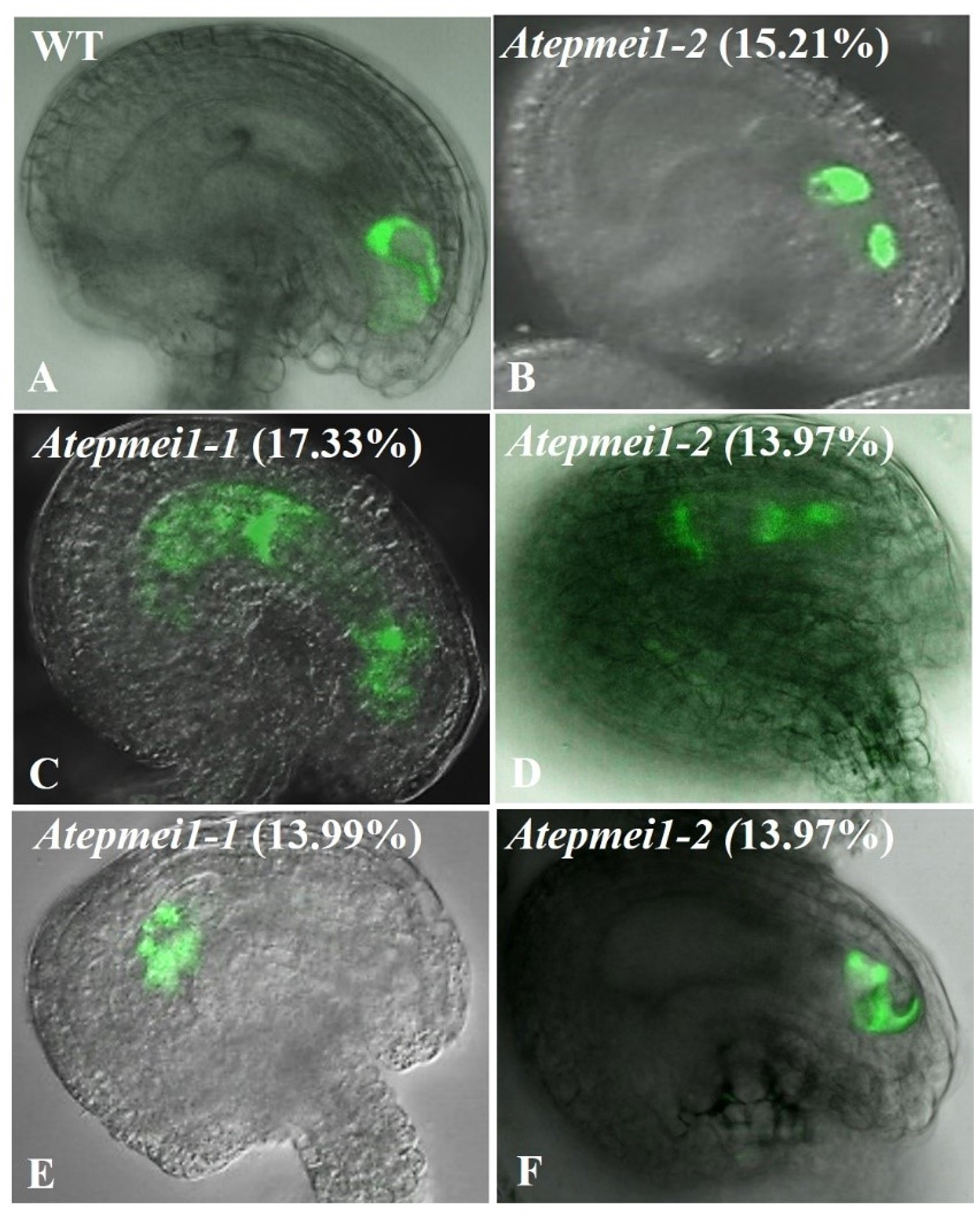
Expression pattern of egg cell-specific marker (DD45) in *Atepmei* ovules. Representative expression patterns are depicted. A) DD45 marker expression in WT, B) In *Atepmeil-2* mutant ovule showing GFP at synergid and egg cell positions, C) Mutant ovule showing diffused GFP expression at micropylar and chalazal ends D) Mutant ovule showing GFP expression at three locations i.e., egg cell, central cell, and antipodal cells, E) GFP expression at the chalazal end, at antipodals position. F) GFP expression at the micropylar end, at the synergids position. Scale bar: 20 pm. Values in parenthesis refer to the frequency of mutant ovules showing indicated GFP expression.

In WT, synergid cell-specific marker (DD31::GFP) lighted up as two dots at the micropylar end (Figure S4). In mutants, GFP was not detectable in 46-54% of embryo sacs (Table S1) while ∼28% of embryo sacs displayed two GFP spots at the micropylar end as found in WT. Besides, some of the mutant embryo sacs showed abnormal GFP expression at locations other than the micropylar end. In 9-16% of ovules, a single GFP spot was detected at the micropylar end (Figure S4 B and Table S2). In 6.89 – 8.26% of ovules, GFP fluorescence was observed at two positions, one at the micropylar end and the other at the egg cell position (Figure S4 C and D), GFP signal at three positions in a row at micropylar end (Figure S4 E) and diffused pattern of GFP localization was observed in 1.72% embryo sacs (Figure S4 F).

Expression of central-cell-specific marker DD65::GFP was also analyzed in both *Atepmei1-1* and *Atepmei1-2* mutants. In WT, this marker expresses in the central region after the fusion of two polar nuclei (Figure S5 A). In *Atepmei1-1* and *Atepmei1-2* mutants, no GFP fluorescence was observed in 45-61% of ovules (Table S2), whereas, 17-23% of ovules showed WT-like normal GFP fluorescence. In 10.08-13.99% of mutant ovules GFP localization was observed away from the normal central position (Figure S5 B and F). In 10.3% of ovules, diffused GFP fluorescence was observed in the entire embryo sac (Figure S5 D). In 10 – 13.99% of ovules, GFP expression was detected at the micropylar end (Figure S5 C and E). Antipodal-cell-specific marker DD1::GFP showed GFP fluorescence as three dots at the chalazal end in WT (Figure S6 A) whereas complete absence of fluorescence was observed in 27-32% ovules in both *Atepmei1-1* and *Atepmei1-2* mutants (Table S2). However, the antipodal-specific marker (DD1) exhibited normal expression pattern in most of the ovules from both the mutant lines. Apart from this, a small percentage of the ovules displayed the GFP expression of DD1::GFP differently. In some ovules (1-5%), diffused GFP fluorescence was observed (Figure S6 B and C), while 10% ovules showed fluorescence at a single position (Figure S6 D). Thus, based on GFP expression data, we conclude that mutation in *AtEPMEI* altered cell specification within the embryo sac (∼50%) and disturbed the location-specific cell identity in the female gametophyte.

Analysis of egg cell marker (DD45) expression in the *pmei* mutant showed GFP at three locations i.e., either in the synergid cell position or in the central cell position along with the egg cell position in 47% of mutant ovules (Figure 6, B,C,D). Similarly, the synergid-specific marker (DD31) expression was also localized to the egg cell position in 20% of mutant ovules, along with its localization at one position in the micropylar end (Figure S4; C,D,E,F). The percentage of ovules expressing synergid specific marker in one cell at the micropylar end and percentage of ovules expressing egg cell marker in one of the synergid cell areas, along with normal egg cell position coincided with the percentage of twin embryos (∼14%) obtained. On the basis of these results, it can be concluded that the additional egg cell(s) are derived either from mis-specified synergids that might still retain the characteristic position at the micropylar region or from mis-specified central cell. This misspecification of gametes appears to result from defects in embryo sac cellularization. Taken together, the above results indicate that *AtEPMEI-*mediated embryo sac cellularization is necessary to prevent synergids and central cells from taking on egg cell fate and cell fate is not determined solely by its location within the embryo sac. It is, however, not clear whether the auxin gradient-mediated gamete specification still holds true in these mutants.

To examine the auxin gradient in *Atepmei*, transgenic *Atepmei1-2* plants carrying DR5::GUS auxin reporter construct were generated. In WT the DR5-driven GUS signal is seen in the nucellus, outside the developing embryo sac at the FG1 stage (Figure 7A). At the FG3 stage, the signal is detectable at the micropylar pole inside the embryo sac (Figure 7B). A strong signal was localized to the micropylar end of the developing embryo sac (Figure 7C) by stage FG4 (after the second mitotic division). A more or less similar type of GUS signal was localized in normally developing *Atepmei* mutant ovules (Figure 7D, E, F), whereas no GUS signal was detected in the abnormally developing ovules (Figure 7G, H, I). These observations suggest absence of strong auxin gradient within the mutant embryo sacs as opposed to WT and WT-like mutant ovules where localized GUS signal is detected. These results indicate altered auxin expression between mutant and WT ovules

**Figure 7.**
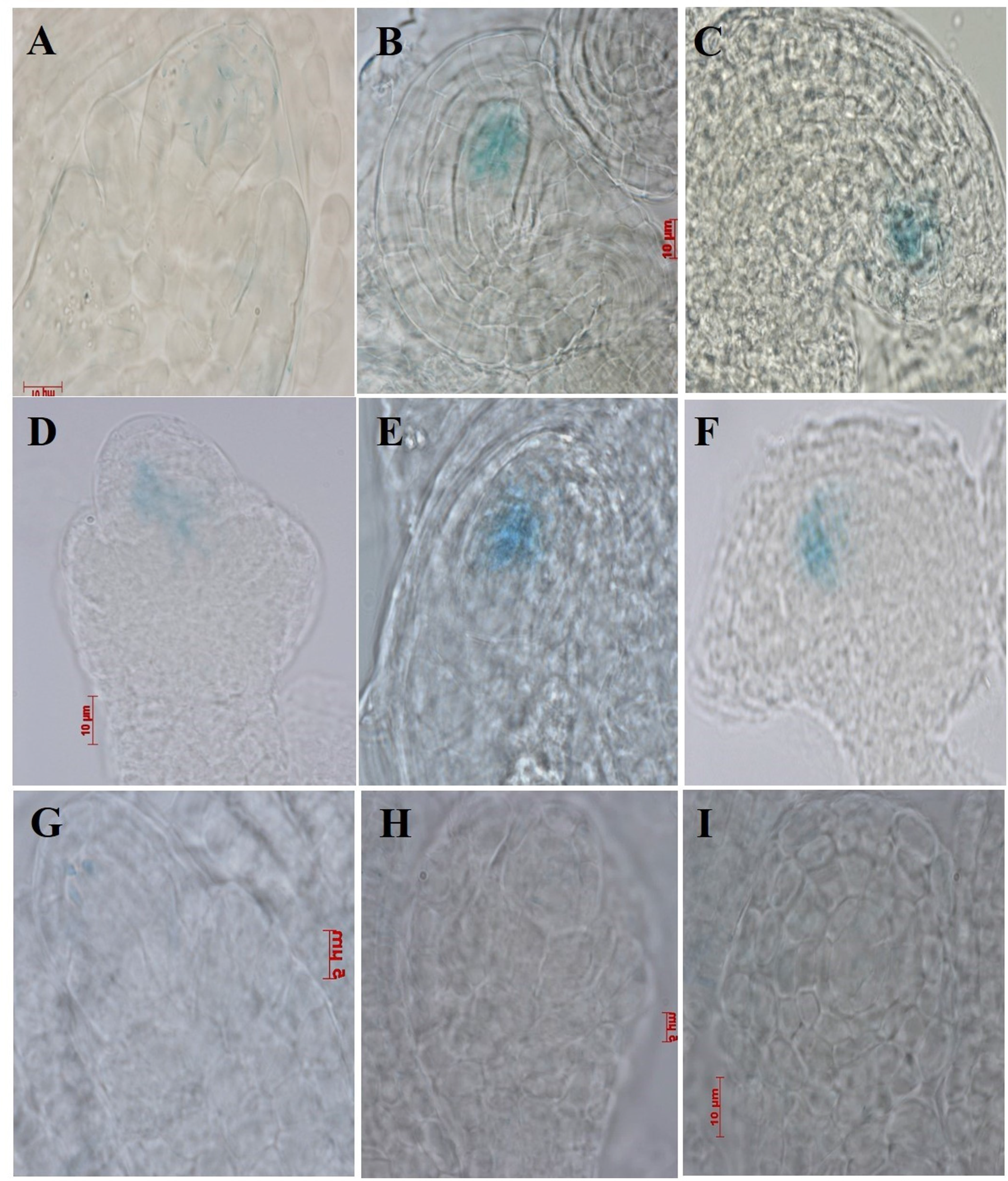
Expression of the synthetic auxin­reporter *DR5::GUS* during female gametophyte development in WT (A-C) and *Atepmeil-2* (D-I). A) At FG1, a faint GUS signal is detected in the embryo sac. B) At FG3 stage, the signal is detectable inside the embryo sac at the micropylar pole. C) Mature ovule showing GUS signal localized near the micropylar end of the embryo sac. D-F) GUS activity in normal-looking embryo sacs at corresponding stages in *Atepmeil-2* mutant. G-I) Little or no GUS activity in abnormal embryo sacs at corresponding stages of female gametophyte development in *Atepmeil-2* mutant.

### Mutation in *AtEPMEI* alters the levels of demethylesterification of cell wall pectin during cellularization of the embryo sac in *Arabidopsis*

Our previous report on *AtEPMEI* promoter analysis (Sharma et al. 2015) confirmed that *AtEPMEI* is spatiotemporally regulated during embryo sac cellularization at the FG5 stage after completion of three rounds of mitosis. During embryo sac cellularization, pectin is loaded in a highly methyl-esterified form. After demethylesterification by the activity of pectin methyl esterase, Ca^2+^ combines with the demethylesterified pectin and forms calcium pectate and becomes rigid. This gives firmness to the cell walls around the gametes and ultimately keeps them in position which helps in the formation of the so-called ‘egg apparatus’ at the micropylar end (Ha *et al*. 1997; Braccini and Perez 2001; Willats *et al*. 2001a; 2001b; Wolf *et al*. 2009; White *et al*. 2014; Wormit and Usadel 2018). We investigated the status of demethylesterification of pectin in cell walls of the embryo sacs of the *Atepmei* mutant using ruthenium red (RuR) staining. RuR stains non-esterified (de-esterified) pectins (Frey-Wyssling, 1976; Western et al., 2001). RuR is a cationic dye having six positive charges and is thereby capable of forming electrostatic bonds with the acidic groups of pectin (Sterling, 1970; Luft, 1971; Lionetti, 2015). Therefore, RuR binding to pectin increases as the degree of methylesterification decreases and vice-versa. Thus, RuR staining could be employed to estimate the extent of de-esterified pectins in WT and *Atepmei* ovules.

WT ovules showed faint pinkish-red staining at the micropylar end of FG4 (Figure 8A) and FG5 stage ovules (Figure 8B), and more intense staining was observed in mature ovules (Figure 8C) followed by very faint staining in post-fertilized ovules (Figure 8D). In contrast, mutant ovules exhibited comparatively more intense staining than WT at all corresponding stages (Figure 8 E-H). More intense staining corresponds to the presence of more de-esterified pectin in the mutant embryo sacs in comparison to WT. This indicates that the normal spatio-temporal process of pectin maturation is disturbed in the *Atepmei1* mutant, consistent with the observed defective cellularization process in the mutant from DIC and cell-specific-marker analyses.

**Figure 8.**
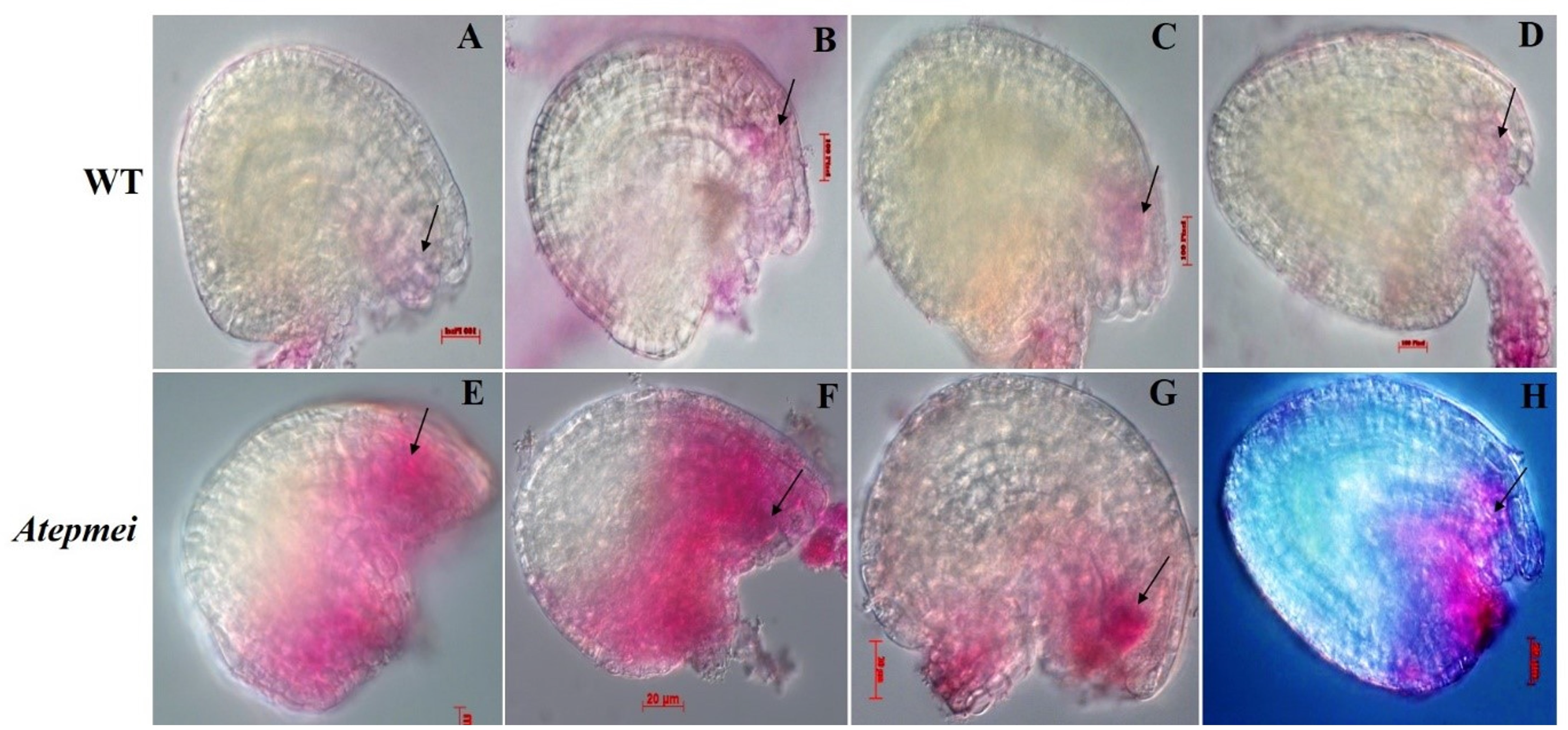
Assessment of de-esterified pectin in the embryo sacs of *Atepmei* mutant ovules by ruthenium red staining. A-D: WT ovule at the FG4 (A), FG5 (B), FG7 (C) and post-fertilization stage (D) showing faint RuR staining near the micropylar end. E-H) *Atepmeil-1* ovules at the FG4 (E), FG5 (F). FG7 (G) and post-fertilization (H) stage showing intense RuR staining. Staining region marked with arrows.

Attempts were also made to assess the status of Ca^2+^-linked pectin i.e., calcium pectate formation in the mutant embryo sacs using tannic acid-ferric-chloride staining, which stains them in black or blue-black (Hornatowska, 2005). Embryo sacs from WT ovules showed faint blackish staining at the micropylar end at FG4 (Figure 9A) and FG5 stages (Figure 9B) and strong staining in mature ovules (Figure 9C), followed by weak staining in post-fertilized ovules (Figure 9D). Embryo sacs from the mutant ovules exhibited comparatively strong staining (Figure 9E-H) compared to WT at all the developing stages, indicating abnormal accumulation of calcium pectin gel in the mutant embryo sacs. In some cases, intense staining was observed throughout the embryo sac. Similarly, post-fertilization ovules from the mutant also showed enhanced staining in comparison to WT. These results further suggest the involvement of the *AtEPMEI* gene in the regulation of PME activity during the cellularization of pre-fertilized ovules and also during the syncytial endosperm development stage.

**Figure 9.**
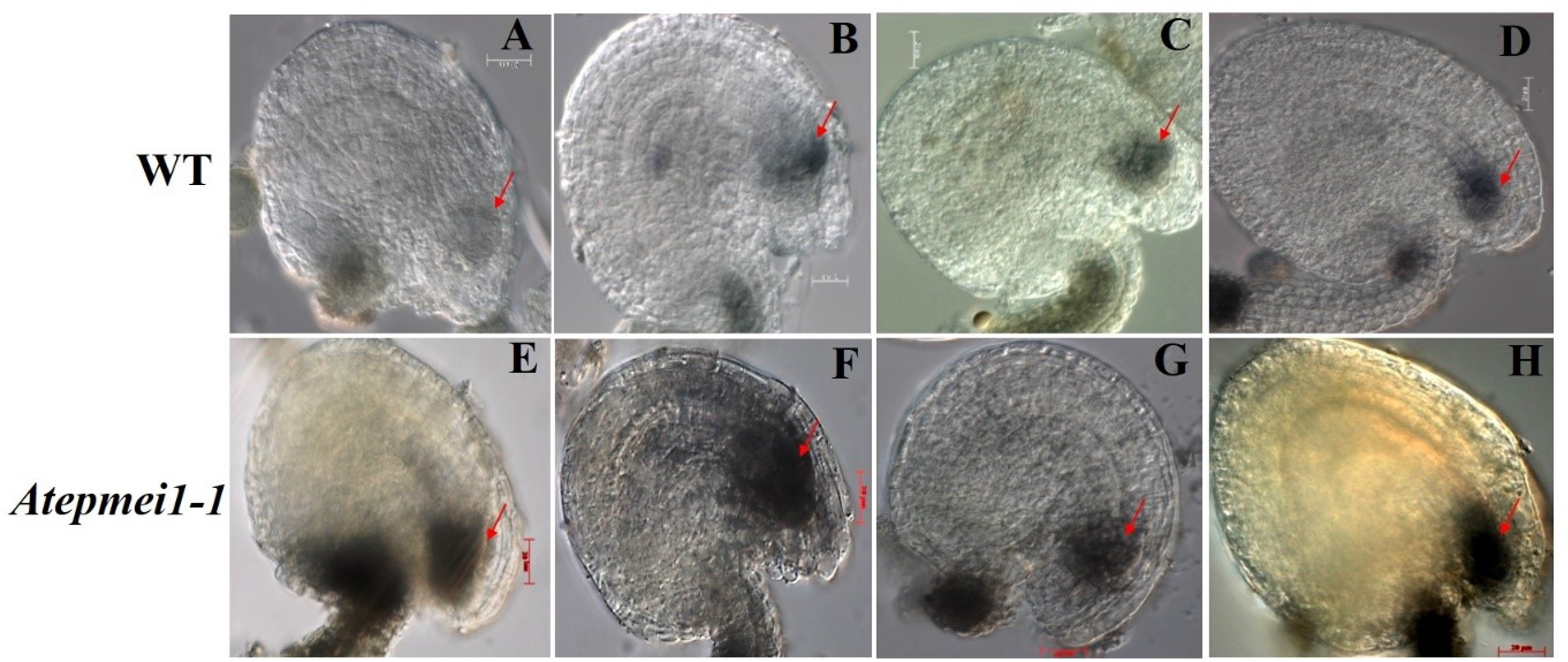
Assessment of calcium pectate accumulation in the embryo sacs of *Atepmei* mutant ovules by tannic acid-ferric chloride staining. A-D: WT ovules at FG4 (A), FG5 (B), FG7 (C) and post-fertilization (D) stage displaying faint to deep staining near the micropylar end. E-H: *Atepmei* ovules at the FG4 (E), FG5 (F), FG7 (G) and post-fertilization. (H) Stage showing intense staining near the micropylar end. Stained regions are marked with red arrows.

## Discussion

The embryo sac of angiosperms is a heterogeneous cell with two distinct poles (micropyle and chalaza). Our understanding of differentiation and development of female gametophyte in angiosperms is still elementary. The nuclei locations in syncytial embryo sac just before cellularization appear to be associated with cell fate of the female gametophyte. Still, there is no proof for the same as the eight nuclei prior to cellularization cannot be readily distinguished (except for size). The gametophyte-cell-specific markers start expression only after cellularization (Susaki et al. 2021). Results of Pagnussat et al. (2009) indicated auxin gradient within the embryo sac prior to cellularisation as the main determinant of location-based cell fate determination. However, a subsequent study (Lituiev et al. 2013) failed to validate it and, instead, suggested that sporophytic auxin influx might contribute to auxin gradient. Studies of various gene mutations belonging to different categories such as cytokinin receptor related (*CKI*), inner centromere protein (*WYRD*), *AMP1*, homeodomain (*eostre*), transcription factors (B3, MYB64, 98, 119, RKD), RNA splicing factors (*CLO, ATO, LIS* etc.,) have clearly indicated that cell fate is not inflexible and gametophyte cells can interchange cell fate despite their location. Study of *lachesis* mutant has shown that egg cell negatively regulates synergid cells from acquiring egg cell fate thereby underscoring the importance of cell-cell communication for maintenance of cell fate (Grob-Hardt et al. 2007, Volz et al. 2012). Similarly, critical role of cell-cell communication in maintaining cell identity is demonstrated in other studies (Kagi et al. 2010; Wu et al. 2012; Tekleyohas et al. 2017). Thus, cell fate not only depends upon the initial acquisition of its identity but also on its subsequent maintenance and reinforcement. Cell walls not only provide structural backbone but also serve as anchor to various enzymes, receptors/sensors, pumps and gates, and regulate movement of substrates and facilitate cell-cell communication. Transcriptome studies of developing female gametophytes have identified differentially expressed genes related to cell wall metabolism including *Atepmei* (Yu et al. 2005; 2007; Steffen et al. 2017), but so far, no gene belonging to this category has been identified to affect female gametophyte cell fate. This is the first report demonstrating that a gene regulating cell wall composition, namely, *Atepmei*, affects cell fate determination in the embryo sac.

*Atepmei* mutants showed high female sterility but male fertility and transmission were unaffected. Given that *Atepmei* promoter is specifically expressed in the embryo sac (Sharma et al., 2015), its loss-of-function is expected to manifest in the female side only. However, loss of *Atepmei* function was not lethal and homozygous mutants could be maintained and perpetuated. This is understandable as *Atepmei* mutation alters specification of gametophyte cells but some embryo sacs still retain essential gametic and accessory cells for seed production. *Atepmei* mutant embryo sacs produced supernumerary functional egg cells that gave rise to twin seeds. Further, developing embryos seen at locations away from the micropylar region indicated that cells acquiring egg cell identity but located away from the micropyle can attract and fuse with male gamete to give rise to the zygote. The mutant phenotype could be fully rescued by constitutive expression of *Atepmei* cDNA confirming that the observed phenotypes are due to loss of function of this gene. *Atepmei* is hypothesized to code for PECTIN METHYLESTERASE INHIBITOR which associates with PECTIN DEMETHYLESTERASE to block its function thereby regulate cell wall dynamics. Hence, loss of *Atepmei* function is expected to alter cell wall composition affecting cell-cell communication including metabolite movement. Thus, the observed altered phenotypes of *Atepmei* mutant ovules/embryo sacs can be linked to disruption of cell-cell communication among various cells of the gametophyte.

Altered cell wall composition was also evident from Ruthenium red and tannic acid-ferric chloride staining of developing ovules. Mutant ovules showed staining over a much broader area than the wild type. The role of cell wall composition for megaspore development is illustrated from callose deposition in megaspore mother cell prior to meiosis and its disappearance after meiosis. This is believed to insulate the megaspore mother cell from surrounding sporophytic cells. PMEIs regulate PMEs, which catalyze the demethylesterification of GalA (α-1, 4-linked D-galacturonic acid) residues of homogalacturonan (HG) in plant cell walls in either block-wise or random manner. This consequently lowers the degree of methylesterification and alters the pattern of methylesterification, which ultimately affects the biomechanical properties of cell walls (Micheli, 2001; Wormit and Usadel, 2018; Shi et al., 2018). During block-wise demethylesterification several consecutive GalA are de-methylesterified. Negatively charged carboxyl groups thus formed, in the presence of Ca^2+^, form “egg-box” structures between the HG molecules, which underlie the formation of pectin gels contributing cell wall strengthening (Wolf et al., 2009; Wormit and Usadel, 2018; Braccini and Perez, 2001). This calcium cross-linked HG increases the amount of bound water, thus maintaining cell wall hydration which in turn affects the rigidity of cell wall (White et al., 2014; Ha et al., 1997; Wormit and Usadel, 2018). Therefore, the involvement of specific PME-PMEI interactions in proper cell wall formation and maintenance around embryo sac nuclei can be inferred. The consequences of defects in embryo sac cellularization manifest as abnormal and mis-positioned gametes with altered cell specification. Cell-type-specific GFP marker line expression studies in the mutant embryo sacs also documented defects in gamete and accessory cell identities. Egg cell-specific marker expression analysis confirmed that in *Atepmei* ovules, more than one egg cell was specified due to the loss of its proper specification mechanism, which resulted in impaired fertilization and subsequent defects in endosperm development.

Based on the present results, we present a schematic diagram to explain how *Atepmei*-mediated regulation of cell wall pectin maturation affects cellularization of the embryo sac in *Arabidopsis* and thereby leads to altered cell fate (Figure 10).

**Figure 10.**
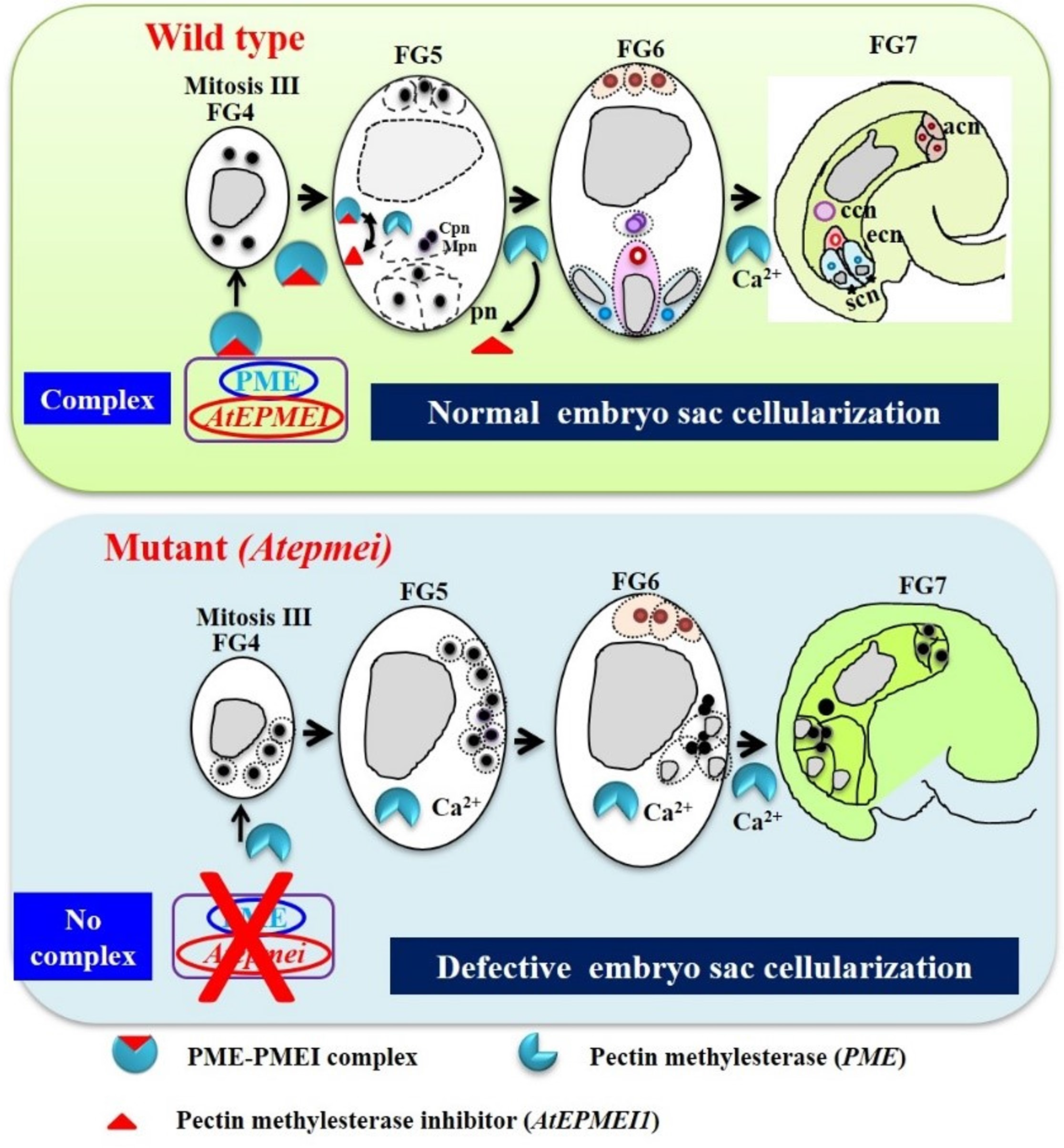
Schematic diagram explaining the probable role of *AtEPMEI*-mediated regulation of cell wall pectin maturation during embryo sac cellularization in *Arabidopsis* based on phenotypic and cell-specific marker expression data. The effect of *AtEPMEI* disruption on female gametophyte development is shown in comparison with WT. In WT, *AtEPMEI* binds to its PME rendering it inactive throughout female gametophyte development with very little cell wall rigidification. In contrast, in mutant ovules, in the absence of PMEI, PME is functional and active leading to demethylesterification of homogalturons (HGs). Subsequent binding of HGs with Ca^2+^ forms ‘egg-box’-like structure causing rigidification of pectin (pectin maturation). Thus, in mutants rigid cell wall forms around the nuclei before reaching the respective positions probably i.e., micropylar, chalazal end, etc. The rigid cell walls of the mutant ovule most likely affect cell-cell communication by preventing flow of metabolites and non-cell-autonomous signals among gametophyte and sporophyte cells. Thus, altered cell-cell communication due to defective cell wall rigidification alters cell fate in mutant female gametophytes. Pn: polar nuclei; ccn: central cell nuclei; ecn: egg cell nuclei; and sen: synergid cell nuclei. Vacuoles are coloured grey. Cell wall formation indicated by dotted lines.

Our promoter analysis study (Sharma et al., 2015) showed that *AtEPMEI* is expressed in embryo sac cells from the early stage till FG8 stage and beyond. Thus, in WT, *AtEPMEI* binds to its PME and keeps check on pectin maturation cell wall rigidification. On the other hand, in the mutant, where AtEPMEI is not produced, PME becomes active causing demethylesterification of HGs. Subsequent binding of Ca^2+^ to demethylesterified HGs causes rigidification of cell walls. Thus, in mutants, PME becomes active even before the completion of three rounds of mitosis leading to formation of rigid cell walls around the nuclei before they occupy their respective positions i.e., micropylar, chalazal ends (Fig. 2B, F, G and H). Possibly, this defect in the cellularization process, also interferes with cell-cell communication, which is known to be crucial for cell fate distribution. In particular, the rigid cell walls of the mutant embryo sacs might interfere with movement of non-cell autonomous signals including auxins among different cells of the female gametophyte and the adjoining sporophyte. The importance of precise regulation of pectin maturation is best illustrated in localized cell wall modifications at the growing tip of pollen grains (Rockel et al., 2008).

Consistent with previous transcriptomics data (Yu et al., 2005; Johnston et al., 2007; Steffen et al., 2007), our findings confirm that *Atepmei* is a major reproduction-related critical embryo sac expressed gene. Further studies on intercellular trafficking of molecules in mutant and WT female gametophytes are needed to unravel the mechanism of PME-PMEI regulated cell fate determination.

## Materials and Methods

### Plant material

Development and screening of transferred DNA (T-DNA) promoter trap population and identification of T-DNA flanking sequences of mutant *A. thaliana* (ecotype Columbia) have been described earlier (**Pratibha** et al. 2013; 2017). Seeds were stored at 4 ^°^C for further use. The seeds of egg, antipodal and synergid cell-specific marker lines (DD45::GFP, DD1::GFP and DD31::GFP) used in the present study were kindly provided by Dr. Gary N. Drews, University of Utah, Salt Lake City, Utah, United States of America. The central cell-specific marker DD65:GFP was obtained from G. Pagnussat’s laboratory. T-DNA mutant line (SALK_063553.52.65.x) was procured from ABRC (Arabidopsis Biological Resource Centre). Egg-cell-specific marker (DD45::GFP) (ET1119) and synergid-cell-specific marker (DD1::GFP) (ET884) lines were obtained from Ueli Grossniklaus’ lab and were analyzed according to Kirioukhova (2011). Central-cell-specific marker was analyzed as per Steffen et al (2007) and Leshem et al (2012).

### Generation of cell specific marker lines in *Atepmei1* mutant background

The marker lines were individually crossed with the *Atepmei1-1* or *Atepmei1-2* mutant lines and F_1_ plants carrying mutant *Atepmei* allele and the GFP marker gene were identified through PCR. Ovules of such F_1_ plants were analysed for marker gene expression.

### Phenotypic characterization of ovules and anthers

Inflorescences from WT and *Atepmei* and complemented plants were kept in 9 parts ethanol: 1-part acetic acid followed by vaccum infiltration for 10-15 min to allow penetration of fixative. Fixation was done overnight at room temperature. Tissue was washed twice with 90% ethanol for 30 min each and kept in 70% ethanol. Fixed pistils from unopened flowers were dissected on a glass slide under Stereo zoom microscope SMZ1500 (Nikon, USA) to release ovules in a drop of chloral hydrate solution. The cleared ovules were then visualized after 5 h, imaged and documented under a microscope equipped with DIC optics (AXIO imager. M1, Carl-Zeiss, GmbH, Germany). Mutant ovules were investigated for nuclei number and their spatial placement within the mature embryo sac in comparison to WT. For further confirmation, buds were emasculated from WT, *Atepmei1-1*, *Atepmei1-2* and observed 2 DAE (days-after-emasculation) for any defects in female gametophyte development.

To study the pollen morphology, fixed flowers were dissected in a drop of chloral hydrate solution under Stereo zoom microscope SMZ1500 (Nikon, USA) to separate anthers from flowers. Cleared anther was imaged and documented under a microscope equipped with differential interference contrast (DIC) optics (AXIO imager. M1. Carl Zeiss, GmbH, Germany).

Pollen viability was determined using Alexander staining (Alexander, 1969). Pollens which are viable stain pinkish while aborted pollen stains green under a light microscope. The working stain solution was diluted 50 times from stock solution. Fresh flowers were dissected under Stereo zoom microscope SMZ1500 (Nikon, USA) to remove anthers on a drop of Alexander’s stain (∼50 µl). Anthers were pressed with pinhead slightly to release pollen. The sample was mounted in the same stain and observed under microscope (AXIO imager. M1. Carl Zeiss, GmbH, Germany) after 30 minutes.

Histochemical GUS assay was performed as described by Jefferson et al (1987) using the chromogenic substrate XGluc (5-bromo-4-chloro-3-indolylglucuronide) (Biosynth). Tissue was incubated in GUS staining solution, vacuum infiltrated for 10 min and left in the staining solution overnight at 37 °C. GUS-stained tissues were then washed with 70 % ethanol twice to remove the chlorophyll and cleared in the chloral hydrate solution (chloral hydrate:water:glycerol, 8:2:1) for at least 5 h. Cleared tissues were observed directly under a microscope (Axio Imager.M1; Carl Zeiss GmbH, Germany).

### Analysis of embryo sac cell-specific marker expression in the mutant plants

To investigate the expression of embryo sac cell-specific markers in a mutant background, homozygous *Atepmei* mutant plants were crossed with different embryo sac cell-specific marker lines. Each marker line was individually crossed with homozygous *Atepmei* mutant lines. Flowers from F_1_ plants were taken on a glass slide and dissected to release ovules under Stereo zoom microscope SMZ1500 (Nikon, USA) in a drop of distilled water. Coverslip was then placed and ovules were analyzed for GFP expression. GFP expression was documented in the mutant background for all the four marker lines. Images were taken using a Zeiss LSM510 confocal microscope. GFP was excited with an argon laser at a wavelength of 488 nm, and emission was detected at 510 nm. In the case of mutant lines, ovules were categorized into different types: as ovule showing normal GFP fluorescence, absence of any fluorescence, fluorescence, at other than normal positions and quantitative data was recorded.

### *In-situ* detection and imaging of de-esterified pectin and calcium pectate (Ca^2+^**-**linked pectin) in ovules

WT and mutant pistils from the flowers at different developmental stages (unopened buds, unopened flowers, open flowers) were selected for pectin localization in the ovules. For localization of de-esterified pectin ruthenium red staining (RuR) was done as described by Hornatowska (2006) and Aggarwal (2009). Pistils were dissected on a glass slide and ovules were gently removed with needles and stained in 100 µl of 0.05% aqueous ruthenium red stain for 3 minutes. The staining reaction was stopped by adding distilled water. Ovules were then observed and the difference in staining intensity was documented in both the mutant and WT by DIC microscopy (AXIO imager, M1, Carl-Zeiss, GmbH, Germany).

Tannic acid-ferric chloride staining was done to stain calcium pectate which stains blue-black (Hornatowska 2006). Plant material was placed in 1% aqueous solution of tannic acid for 10 minutes. After that 3% aqueous solution of ferric chloride was added to the sample and incubated for 3 min and the staining reaction was stopped by adding distilled water. The difference in staining intensity between WT and mutant was observed under a DIC microscope (AXIO imager, M1, Carl-Zeiss, GmbH, Germany).

## Acknowledgements

This work was supported by the Council of Scientific and Industrial Research (project nos. MLP-072 and BSC-0107: at CSIR-IHBT, Palampur and MLP-102 at CSIR-CCMB, Hyderabad) as well as senior research fellowship to I.S. and the Indian Council of Agricultural Research through the National Agricultural Innovation Project (Grant No. NAIP-4157). Y.S. also acknowledge Institute of Eminence – University of Hyderabad (IoE-UoH) research grant (UoH-IoE-RC3-21-009). P.M. acknowledges UGC for her Kothari Postdoctoral Fellowship.

## Authors contributions

I.S. and P.M. conducted the experiments. R.S., S.R.B. and Y.S. mutant generation, designed the experiments for the analysis of the mutant, compiled the data and wrote the paper.

